# Inference of Presynaptic Connectivity from Temporally Blurry Spike Trains by Supervised Learning

**DOI:** 10.1101/2022.10.20.513050

**Authors:** Adam D. Vareberg, Jenna Eizadi, Xiaoxuan Ren, Aviad Hai

**Affiliations:** Department of Biomedical Engineering, University of Wisconsin–Madison; Department of Electrical and Computer Engineering, University of Wisconsin–Madison

## Abstract

Reconstruction of neural network connectivity is a central focus of neuroscience. The ability to use neuronal connection information to predict activity at single unit resolution and decipher its effect on whole systems can provide critical information about behavior and cognitive processing. Neuronal sensing modalities come in varying forms, but there is yet to exist a modality that can deliver readouts that sufficiently address the spatiotemporal constraints of biological nervous systems. This necessitates supplementary approaches that rely on mathematical models to mitigate physical limitations and decode network features. Here, we introduce a simple proof-of-concept model that addresses temporal constraints by reconstructing presynaptic connections from temporally blurry data. We use a variation of the perceptron algorithm to process firing rate information at multiple time constraints for a heterogenous feed-forward network of excitatory, inhibitory, and unconnected presynaptic units. We evaluate the performance of the algorithm under these conditions and determine the optimal learning rate, firing rate, and the ability to reconstruct single unit spikes for a given degree of temporal blur. We then test our method on a physiologically relevant configuration by sampling network subpopulations of leaky integrate-and-fire neuronal models displaying bursting firing patterns and find comparable learning rates for optimized reconstruction of network connectivity. Our method provides a recipe for reverse engineering neural networks based on limited data quality that can be extended to more complicated readouts and connectivity distributions relevant to multiple brain circuits.

## 1. INTRODUCTION

There remain fundamental difficulties in decoding brain activity and relating it to function across multiple spatiotemporal and sensitivity scales (1–3). Nervous systems operate as composite biological networks critical to diverse functions including motor tasks, sensory perception, sensation, and memory, carrying intricacies at the molecular level across expansive interconnected units and circuits (4–6). Consequently, the interpretation of experimental neural data that are limited in scope relies heavily on innovative analysis and automated learning methods to extract and predict network features relevant to function (7,8). Such methods have been used to decipher neural circuitry directly from data and predict brain function in clinical context, unlocking new possibilities for treatment of neurological diseases (9,10). Extracting meaningful information depends on data quality, scope and type. Steady progress is made with upgrading sensing and hardware capabilities to improve the selection and scale of neurobiological data available to *in silico* studies. High density neural probes with increased spatial coverage provide chronic electrophysiological readouts of direct spiking activity from hundreds and thousands of units (11–14). Other readout types with reduced temporal detail such as optical imaging of slow calcium fluctuations can still offer large scale single unit activity critical to understanding neural circuitry (15–17) and can be deconvolved computationally to uncover underlying action potential information with reasonable precision (18,19). Noninvasive approaches present less tractable options for deconvolution both temporally and spatially (20–22). Whole-brain neuroimaging of blood flow was found to partially correlate with electrophysiology (23) and neurotransmitter systems (24,25), but emerging sensors are now providing direct volumetric recordings of calcium levels (26,27) neurotransmitter dynamics (28–30) and electromagnetic fields (31,32). The temporal resolution for these techniques is limited depending on the phenomenon studied and the physical hardware limitations of the modality in use. Developing new computational strategies to mitigate these challenges by optimizing network reconstruction and taking temporal limitations into account—can aid in leveraging new data types for uncovering the electrophysiological underpinning of brain activity.

Different approaches have been demonstrated for processing subsampled and low resolution datasets to infer connectivity (33–37). Some of these methods exploit specialized linear models to enhance information extracted from correlated spiking rates to improve performance for predicting synaptic weights (36). Other studies demonstrate increased performance by assuming distinct nonlinearity types and low noise of temporally restricted data to successfully reconstruct network connections (33,38). Supervised learning of feed-forward perceptron networks are specifically suited for making predictions related to synaptic strength and information storage with missing information and silent synapses (39,40). By matching learning rates with different network states, it is possible to make accurate spike predictions and to quantify information storage and learning capabilities. However, quantifying the ability to predict presynaptic weights and spiking activity based on temporal quality of the original data, and its relation to network activity levels and learning rates, has not been addressed yet and could provide an expanded protocol for reverse engineering neural networks using data limited in scope. In this work, we develop a learning model based on adaptations from the perceptron, which has been used previously to reconstruct presynaptic connectivity from binary events (40). We use presynaptic spike train data with increasing degrees of blurriness as input for a supervised learning, and demonstrate the performance of the algorithm in learning presynaptic weights and predicting postsynaptic spikes. We then expand on these findings by quantifying the effect of temporal resolution on performance at multiple neuronal firing rates to provide a general guideline for optimal learning rates that can be used under each of these conditions. We then test the physiological relevance of the algorithm by implementing a three-layer leaky-integrate-and-fire (LIF) network model displaying bursting activity and evaluate performance with different degrees of access to presynaptic spike data. This work lends to neuroscience a useful technique for reconstructing biological neural networks under temporally constrained sensing conditions relevant to existing brain recording modalities.

## 2. METHODS

### 2.1 Feedforward network initialization

We constructed a network model containing a heterogenous presynaptic cell population (*n* = 200) with simulated spike trains at firing rates ranging between 5 and 80 Hz. Half of the cell population was unconnected and half consisted of excitatory and inhibitory cells (4:1 ratio) connected to a postsynaptic cell, with weights *w_i_* ∈ (0 8] and [-8 0) (mV), *i* ∈ {1..n} distributed uniformly (Fig. 1a). Spike trains were generated over 5000 trials of 1000 ms duration for each presynaptic cell, with each spike representing a single-millisecond event (Fig. 1b). Postsynaptic spike event *y* was determined by a simple integrate-and-fire mechanism:

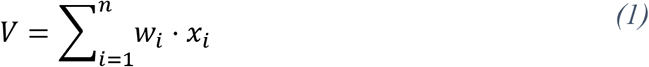

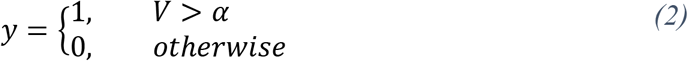

**Figure 1.**
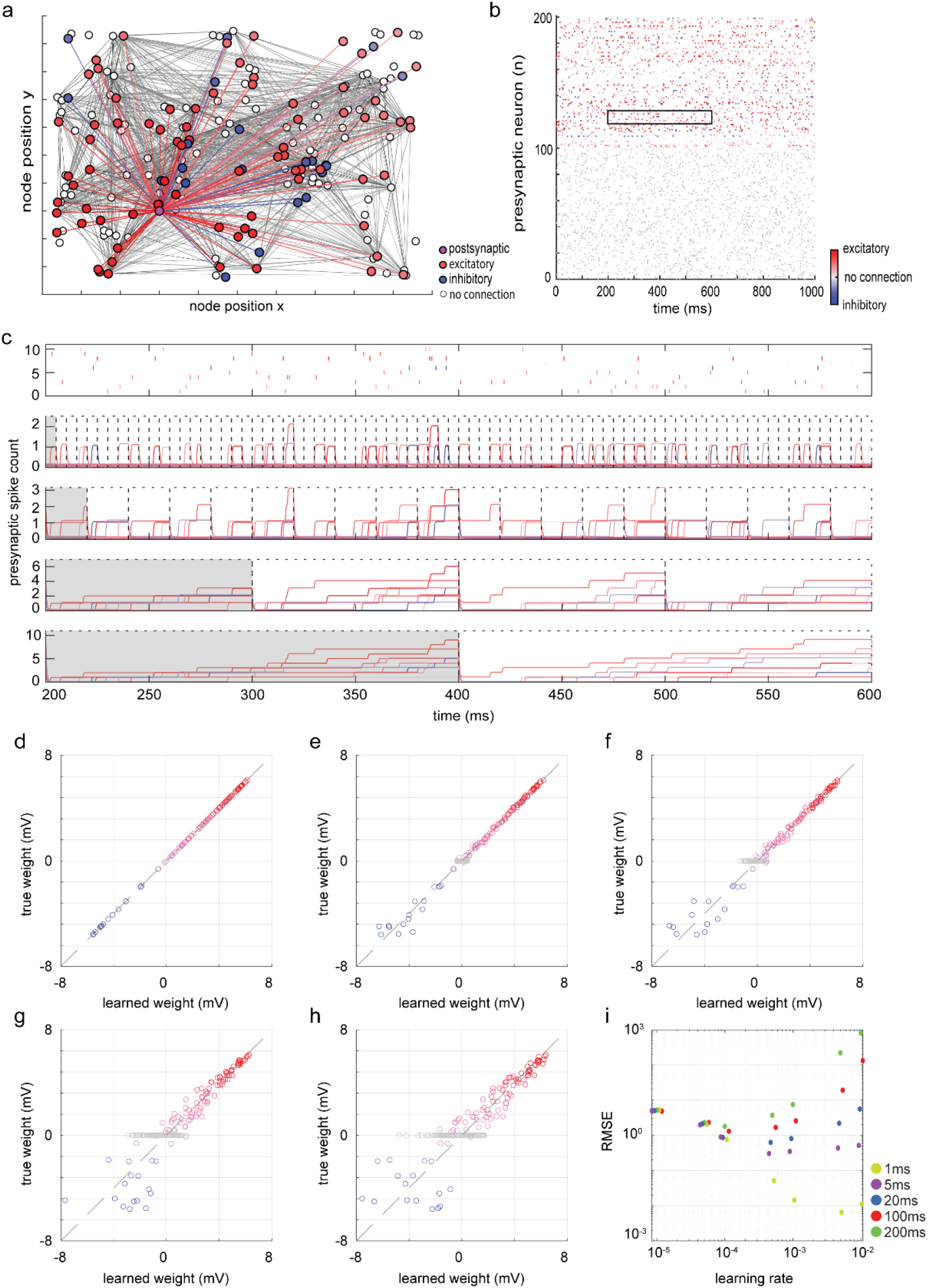
Performance of adapted perceptron on varying presynaptic ‘blurriness’ for an example network. **(a)** Representation of a neural network, with a distribution of inhibitory and excitatory presynaptic cells connected to a single postsynaptic neuron of interest, and neurons with little to no connection to the postsynaptic cell. **(b)** 1000ms of binary spiking data for the presynaptic cells. Highlighted by the black box is a smaller subset of 10 neurons over 400ms duration. **(c)** Depiction of the summation of the presynaptic spike highlighted in panel b into lower resolution time bins. From top to bottom are the discrete 1ms spikes, followed by 5ms, 20ms, 100ms and 200ms bins. Gray boxes depict respective time bin sizes representing the temporal detail fed to the perceptron. **(d-h)** Comparisons of the inferred presynaptic weights with true weights. Each panel depicts the inferred weights provided by the learning rate with the lowest RMSE for each bin size, in order of ascending sizes. **(i)** The root-mean-square error was calculated for the inferred weights against the true weights and plotted for each combination of predetermined bin size and learning rate.

Where *V* is the postsynaptic membrane potential, *α* = 20 mV is the postsynaptic firing threshold, *x_i_* and *w_i_* are the spike input and weight for neuron *i*, respectively, and *n* is total number of presynaptic neurons.

### 2.2 Firing rate perceptron adaptation

We utilized a modified perceptron algorithm for supervised inference of presynaptic connectivity based on firing rate data. A randomized weight matrix was initialized prior to the first iteration. Weights were updated every iteration according to the product of learning rate *lr* and the sum (Δ_*b*_) of each discrete error *δ* = *y* – *y_l_* within time bin *b*:

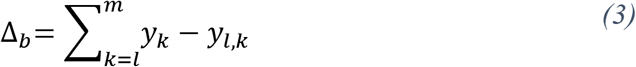

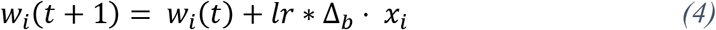

Where *l* and *m* are the first and last discrete timepoints in bin *b*, respectively, and *y* and *y_l_* are the true and predicted postsynaptic responses, respectively. The algorithm was trained over the first 2500 trials for a total of 100 iterations for each dataset or condition. Bin sizes of 1ms, 5ms, 20ms, 100ms and 200ms were investigated. For each bin size, seven learning rates were independently investigated (1·10^-2^, 5·10^-3^, 1·10^-3^, 5·10^-4^, 1·10^-4^, 5·10^-5^, 1·10^-5^), for a total of 35 Resolution-Rate Combinations (RRC). Temporal resolution of data was defined as summation of ground truth single spike events for a given bin size (Fig. 1c) and fed into the modified perceptron as input data by randomly redistributing the calculated sum of spikes to discrete millisecond events within the same bin size.

### 2.3 Inference of synaptic weights

The performance *P* of the adapted perceptron model corresponding to the prediction quality of postsynaptic firing rates, was quantified according to equations *(5)-(7)*:

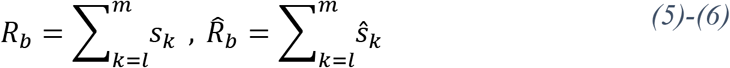

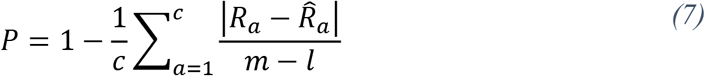

Where, *R_b_* and 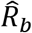 represent the sum of true and predicted discrete postsynaptic spike events *s_k_* and 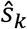, respectively, within bin *b* and averaged over the total number of bins *c* in a training set. We used root-mean-squared error (RMSE) to determine the ability of the adapted perceptron to reconstruct presynaptic weight matrices on the final learned weight matrices to quantify the inferred connectivity provided by the model for each RRC (Fig. 1d-i).

### 2.4 Spike predictions

We evaluated the accuracy of single-spike predictions by generating postsynaptic response from 5000 newly generated 1000 ms trials of presynaptic spike trains. Predicted postsynaptic spikes were calculated using inferred weights and compared to the postsynaptic spike generated using true weights. We determined true positive rate (TPR) for sensitivity and true negative rate (TNR) for specificity, as described in equations *(8)-(9)*, where *S* represents the subset of events in which a postsynaptic spike occurs (*s_p_* = 1) and N represents the subset of events in which there is not a postsynaptic spike:

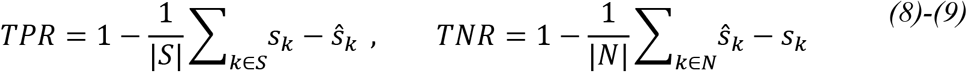

### 2.5 Leaky integrate-and-fire network

A LIF network model was implemented to evaluate performance with physiologically relevant bursting spike patterns. Equation *(10)* describes the membrane potential (*V_m_*) for a given neuron at time *t* + *Δt* as a function of resting potential (*V_e_* = −75 mV), membrane current (*I_m_*), membrane resistance (*R_m_* = 10 MΩ), and membrane capacitance (*C_m_* = 100 pF), where time constant *τ_m_* = *C_m_* × *R_m_* = 1 ms:

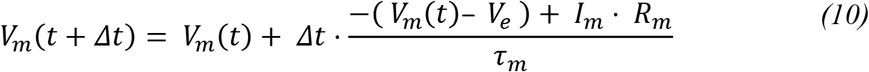

Membrane current is defined as a weighted sum of presynaptic spike defined in equation *(1)* and normalized by the postsynaptic current (*I_post_* = 1 nA). For times *t* where the membrane voltage was greater than the threshold voltage (*V_th_* = –65 mV), an action potential occurred, and membrane voltage was set to 15 mV. For subsequent *t* + *Δt*, neurons hyperpolarized to the reset potential *V_reset_* = –80 mV.

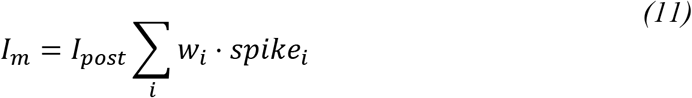

A LIF-based three-layer network was built with a first layer (L1) consisting of spike trains generated using equations *(1)-(2)* at temporal step of 0.2 ms. Resting L1 firing rate was set to 5 Hz, with five 150 Hz bursts, lasting up to 12 ms, injected asynchronously into each second of data. 200 unique L1 neurons were fed into the LIF model to produce voltage traces for each layer 2 (L2) neuron. This process was repeated for 200 L2 neurons acting as presynaptic inputs to layer 3 (L3) to generate leaky postsynaptic data. L2 data was temporally compressed to 1 ms resolution and methods from Section 2.1 were repeated to produce simple integrate-and-fire L3 trains. By constraining learned weights between [-8 8] (mV), four variations of L3 scanerios were tested; unconstrained integrate-and-fire (IFU), unconstrained LIF (LIFU), constrained integrate-and-fire (IFC) and constrained LIF (LIFC). Each network was tested for different fractions of neurons recorded within the network to test performance in common experimental scenarios with limited accessibility (20%, 50%, 80%, 85%, 90% and 95%). In all cases, the same L1 data were used to produce and learn on 5000 s of L3 data.

## 3. RESULTS

### 3.1 Baseline weight inference with increasing presynaptic blurriness

The ability of the adapted perceptron to reconstruct synaptic weights with training sets of varying levels of presynaptic temporal detail is demonstrated in Fig. 1 and Fig. 2. An example network (Fig. 1) was reconstructed based on spike data with increasing temporal bin sizes (Fig. 1d-h, bin sizes: 1, 5, 20, 100 and 200 ms, respectively). We quantified reconstruction accuracy by calculating the minimal RMSE of inferred weights for different bin sizes and learning rates tested (Fig. 1i). Fig. 2 shows iterative prediction accuracy for multiple networks (*n* = 10, 1-100 iterations). The average RMSE over all datasets and RRCs reached minimal values of 6.93·10^-3^ ± 3.02·10^-4^, 0.28 ± 0.02, 0.59 ± 0.03, 1.30 ± 0.04, 1.92 ± 0.09, in order of ascending bin size (Fig. 2f) with respective Pearson correlations of r = 1.00, 1.00, 0.98, 0.92 and 0.85, each (*p* ~ 0), demonstrating that the ability to reconstruct presynaptic weights diminishes with decreased temporal resolution. For small bin sizes (b < 20 ms, Fig. 2a-c) learning converged to maximal performance with fewer iterations at high learning rates. For large bin sizes (b > 20 ms, Fig. 2d-e) high learning rates failed to converge weight predictions. However, we found that with smaller learning rates, performance was improved: RMSE at *lr* = 1·10^-2^ was 3.69·10^2^ ± 81.1 (r = 0.39, *p* ~ 0) and 8.17·10^2^ ± 27.0 (r = 0.71,*p* ~ 0) for bin sizes of 100 and 200 ms, and 4.44 ± 0.63 (r = 0.39, *p* ~ 0) and 8.90 ± 1.15 (r = 0.59, *p* ~ 0) at *lr* = 1·10^-3^, respectively. Performances for smaller learning rates and large bin sizes were not asymptotic at maximum number of iterations tested (100), indicating improved optimum can be reached with higher number of iterations at the expense of algorithm runtime.

**Figure 2.**
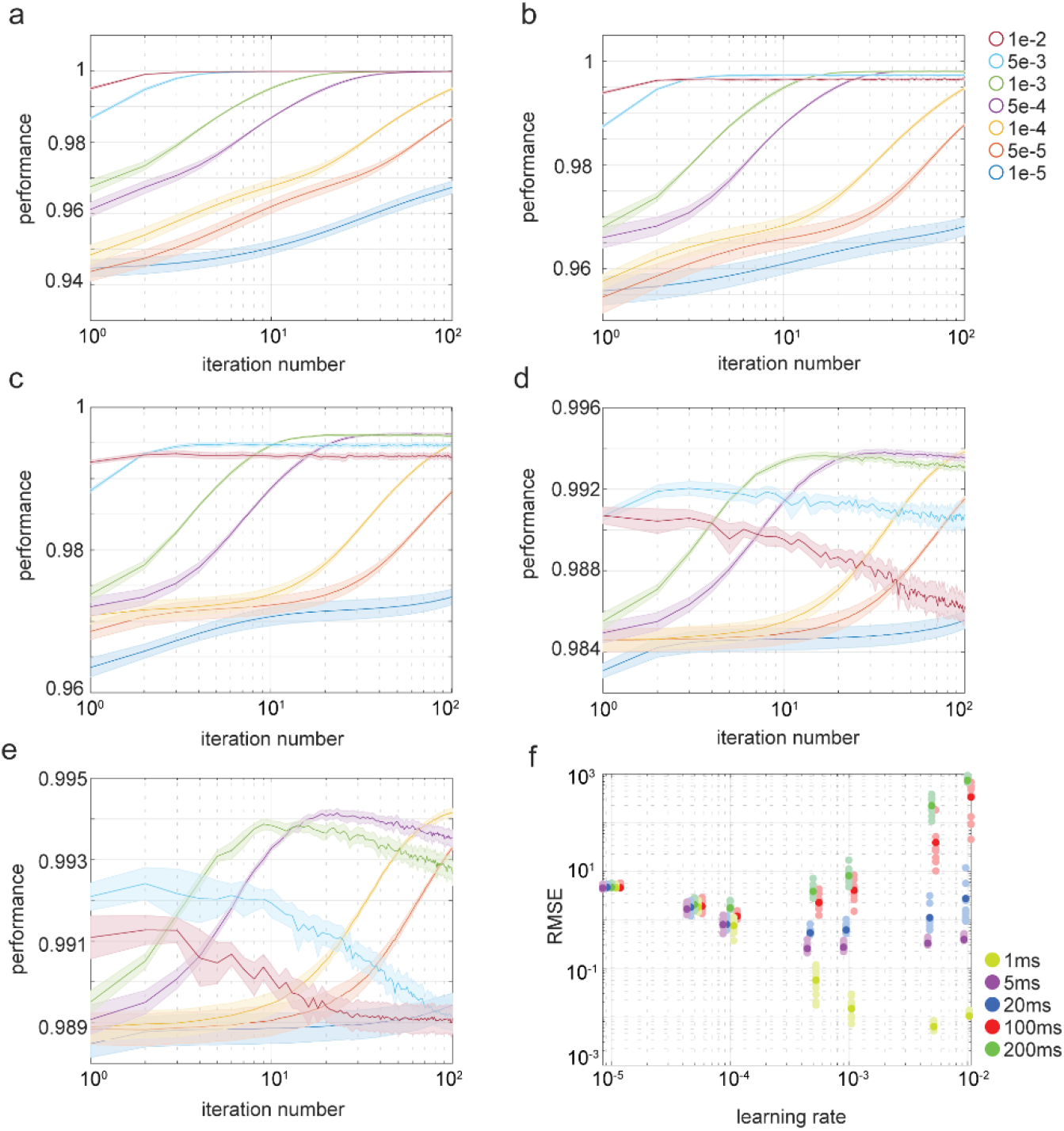
Learning-rate-dependent iterative performance for temporally constrained presynaptic inputs. **(a-e)** Iterative performance of the adapted perceptron. Each panel depicts the performance of the learning rates used in this study for a particular bin size, ordered by bin sizes of 1 ms, 5 ms, 20 ms, 100 ms and 200 ms, respectively. **(f)** Collective RMSE for n = 10 datasets. The average RMSE of these datasets is represented by the darker shade. Mean firing rate was 20 spikes/sec.

### 3.2 Firing rate dependence of weight inference

To find conditions for improved connectivity reconstruction, we turned to comparing firing rate dependent performance for low and high temporal resolution data (Fig. 3). For the lowest temporal bin size tested (200 ms) with mean presynaptic firing rates of 5, 10, 40, and 80 Hz, (Fig. 3a-d), we found minimal average RMSE values of 5.50 ± 0.20 (r = 0.24, *p* ~ 0), 4.50 ± 0.16 (r = 0.54, *p* ~ 0), 0.64 ± 0.05 (r = 0.98, *p* ~ 0), and 0.38 ± 0.02 (r = 0.99, *p* ~ 0), respectively. Lower firing rates (5 Hz and 10 Hz, Fig. 3a-b) exhibited reduced performance instability compared with higher firing rates (40 Hz and 80 Hz, Fig. 3c-d), but low quantity of information for small number of spikes predictably prevented proper convergence of learned weights to true weights (Fig. 3e-f). For higher firing rates, we found greater performance variability and instability, but optimal learning rates utilized the availability of data properly, displaying convergence between learned and true weights (Fig. 3g-h). The mean RMSE for the 200 ms, 5 Hz RRC ranged between 5.50 and 22.8. Comparatively, the respective range for the 80 Hz case ranged between 0.38 and 39.8. Based on these and all other RRCs tested, we obtained a generalized map that defines the optimal learning rate for a given temporal bin size and presynaptic firing rate (Fig. 3m).

**Figure 3.**
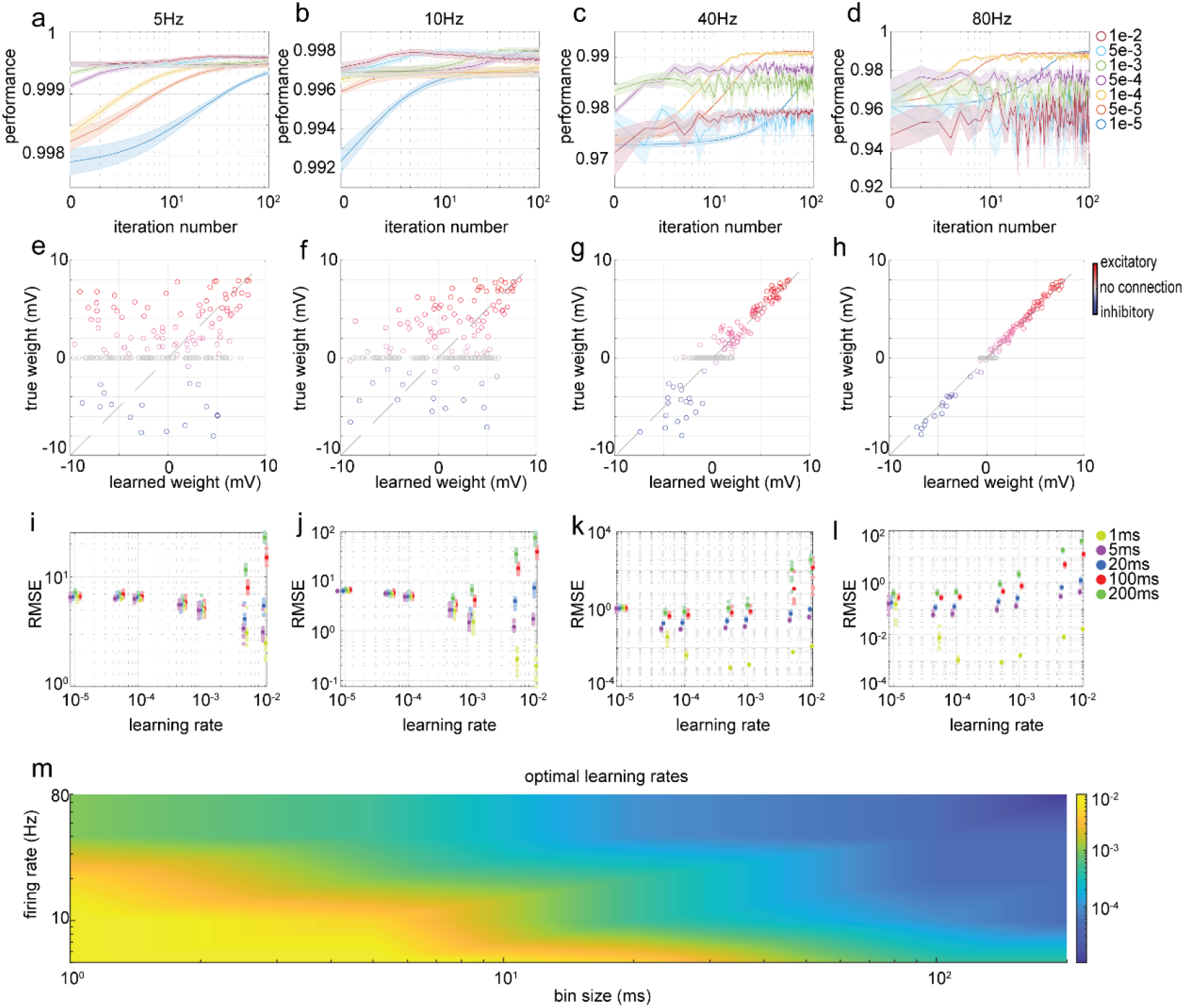
Effect of Presynaptic Firing Rate on Performance of Adapted Perceptron. **(a-d)** Iterative trends on spike redistribution performance for varying presynaptic firing rates. Each panel represents the performance of learning rates as a mean and standard deviation (n=10) of datasets under 200 ms temporal resolution condition. **(e-h)** Comparisons of learned weights against true weights for 200 ms time bins. Each panel represents a sample dataset, selected from the learning rate that provides the lowest average RMSE, as shown in panels i-l. **(i-l)** Collective RMSE for multiple networks (n = 10) for differing presynaptic firing rates, bin sizes, and learning rates, with the average represented by the darker shade. **(m)** A single map was generated to provide optimal learning rates for each RRC.

### 3.3 Single-spikes Predictions

Based on our recipe, we quantified the ability to properly predict activity at the single-spike level (Fig. 4). Figure 4a introduces subsets of predicted spike trains based on weights learned from networks firing at 20 Hz at multiple temporal bin sizes. Predictably, TPR indicated high sensitivity of the algorithm to single-spike predictions for high temporal resolution data that deteriorated as resolution decreased (Fig. 4b). Mean TPR values were 99.85 ± 0.02%, 95.52 ± 0.21%, 90.65 ± 0.40%, 78.64 ± 0.95% and 70.76% ± 1.36%, in order of ascending bin size (Fig. 4b). Specificity of the algorithm persisted as temporal resolution decreased, with mean TNR values of >99.99 ± 3.40·10^-4^ %, 99.91 ± 4.73·10^-3^ %, 99.80 ± 0.01%, 99.56 ± 0.01% and 99.38 ± 0.03% (Fig. 4c). For other presynaptic firing rates, networks with mean spike rate of 40 Hz provided the highest average prediction of true positives in a 200 ms environment, with 13.98 ± 1.85%, 34.81 ± 1.73%, 90.86 ± 0.87% and 80.62 ± 3.68% mean TPR for 5, 10, 40 and 80 Hz, respectively (Fig. 4e). The algorithm remains highly specific under these test conditions, as demonstrated by the respective false positive rates of 98.79 ± 0.16%, 99.35 ± 0.05%, 99.61 ± 0.05% and 99.92 ± 0.02% (Fig. 4f).

**Figure 4.**
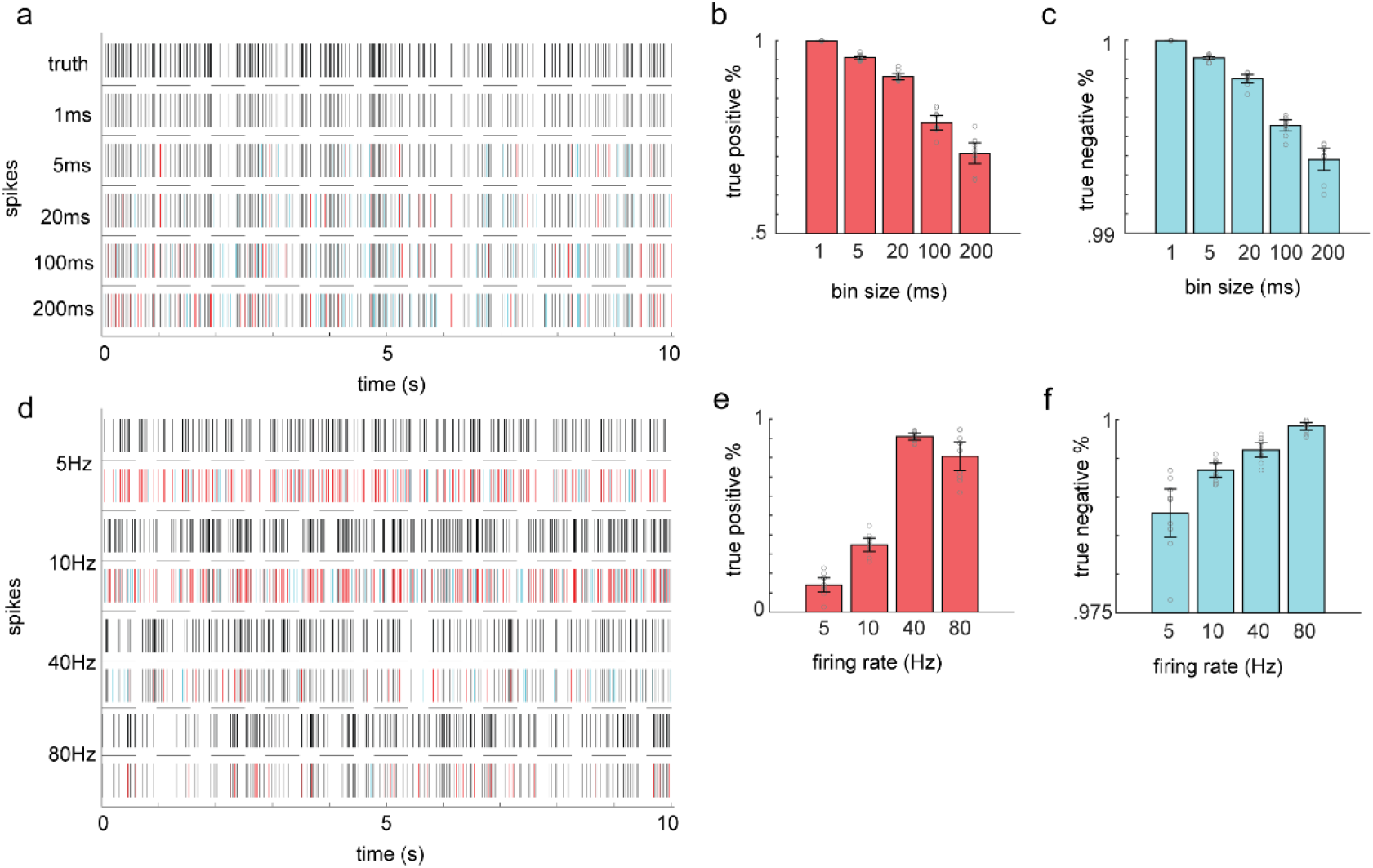
Postsynaptic Spike Prediction from Learned Presynaptic Weights. **(a)** 10 seconds of spike prediction is shown. From top to bottom are the true postsynaptic spikes, followed by the predicted spikes for 1 ms, 5 ms, 20 ms, 100 ms and 200 ms time bins, respectively. Grey spikes indicate correct predictions, while red and cyan represent failed predictions and false positives, respectively. **(b)** Average (n = 10) percentage of correct predictions for each 20Hz time bin. **(c)** Performance against false positives for each 20 Hz time bin, represented as the average (n = 10) percentage of correctly predicted non-spike events. **(d)** Representation of 200ms resolution spike prediction for presynaptic firing rate, with each subsequent two rows demonstrating true spikes and predicted spikes, respectively, for the firing rate. **(e)** Average (n=10) percentage of correct predictions for differing presynaptic firing rates and 200 ms time bins. **(f)** Performance against false positives for differing presynaptic firing rates and 200 ms time bins, represented as the average (n = 10) percentage of correctly predicted non-spike events.

### 3.4 Physiologically relevant Model

Experimentally recorded neuronal activity displays bursting spiking patterns at lognormal firing distributions, and usually provides access to only a fraction of a given network circuit (4). To test performance in physiologically relevant scenarios we implemented a network of LIF neuronal models displaying bursting spiking patterns and tested the ability to reconstruct connectivity with variable degrees of accessibility to presynaptic data and temporal blur (Fig. 5). Stimulation applied through the first layer (L1) manifested in biologically relevant firing bursts at frequencies ranging between 5.83 and 26.96 Hz (*μ* = 16.98, *σ* = 3.57) in L2 neurons (Fig. 5a). Further, we tested performance with learned weights constrained to a physiological range of [-8 8] (mV). Performance in unconstrained integrate-and-fire (IFU), constrained integrate-and-fire (IFC), unconstrained leaky integrate-and-fire (LIFU) and constrained leaky integrate-and-fire (LIFC) configurations was analyzed by determining the RMSE for inferring weights of L2 neurons presynaptic to L3. RMSE values for varying degrees of temporal blur and accessibility to L2 activity data were determined for the respective subset of L2 neurons analyzed (Fig. 5b-e).

For 1 ms temporal resolution IFU, optimal learning rates of 5·10^-4^, 5·10^-3^, and 5·10^-5^ achieved minimum RMSE values of 2.72, 1.09 and 3.31 for 20%, 50% and 100% of the network accessible to the algorithm, respectively. When introducing very high temporal blur (200 ms bins), respective performance values predictably degraded by 13.6 - 62.5% to RMSE values of 5.24, 2.96 and 4.34 with optimal learning rates of 5·10^-5^, 1·10^-3^ and 5·10^-4^. In LIFU cases representing a multilayer LIF network with unconstrained synaptic weights (Fig. 5d), RMSE degraded with increased blur for 20% and 100% of spike data accessible to the algorithm (from 4.04 to 4.89 and 3.84 to 4.02, for 1 ms and 200 ms, respectively) but improved for the 50% case (from 2.91 to 2.40). In addition, 200 ms temporal blur also showed improvement from IFU to LIFU (7.3 - 23.1%). This suggests that low temporal resolution data and may be favored for predicting connectivity in LIF models due to jittery presynaptic input contributing to spike generation compared with naive memoryless feedforward networks (41). Moreover, having access to only a limited fraction of network data can result in higher performance in some cases and with the number of iterations and configuration tested. Constraining synaptic weights to physiological values showed a predictable decrease in performance for all low temporal blur cases and varying degrees of performance change for high temporal blur compared with unconstrained cases (Fig. 5c and 5e). For LIFC (Fig. 5e), 1 ms bins yielded RMSE values of 4.23, 2.98 and 3.70 for 20%, 50% and 100% of the network accessible to the algorithm while increasing temporal blur to 200 ms had varying effect on performance with a large reduction for 20% (28.2%, RMSE = 5.89), moderate improvement for 50% (17.1%, RMSE = 2.54) and minimal change for 100% (2.76%, RMSE = 3.81). Overall, the ability of our method to reconstruct connectivity of bursting LIF networks is less predictable compared with simple IF networks and heavily depends on temporal resolution and the network coverage, demonstrating the necessity for choosing optimal learning parameters to infer weights from a proper subset of presynaptic inputs (Fig. 5f).

**Figure 4.**
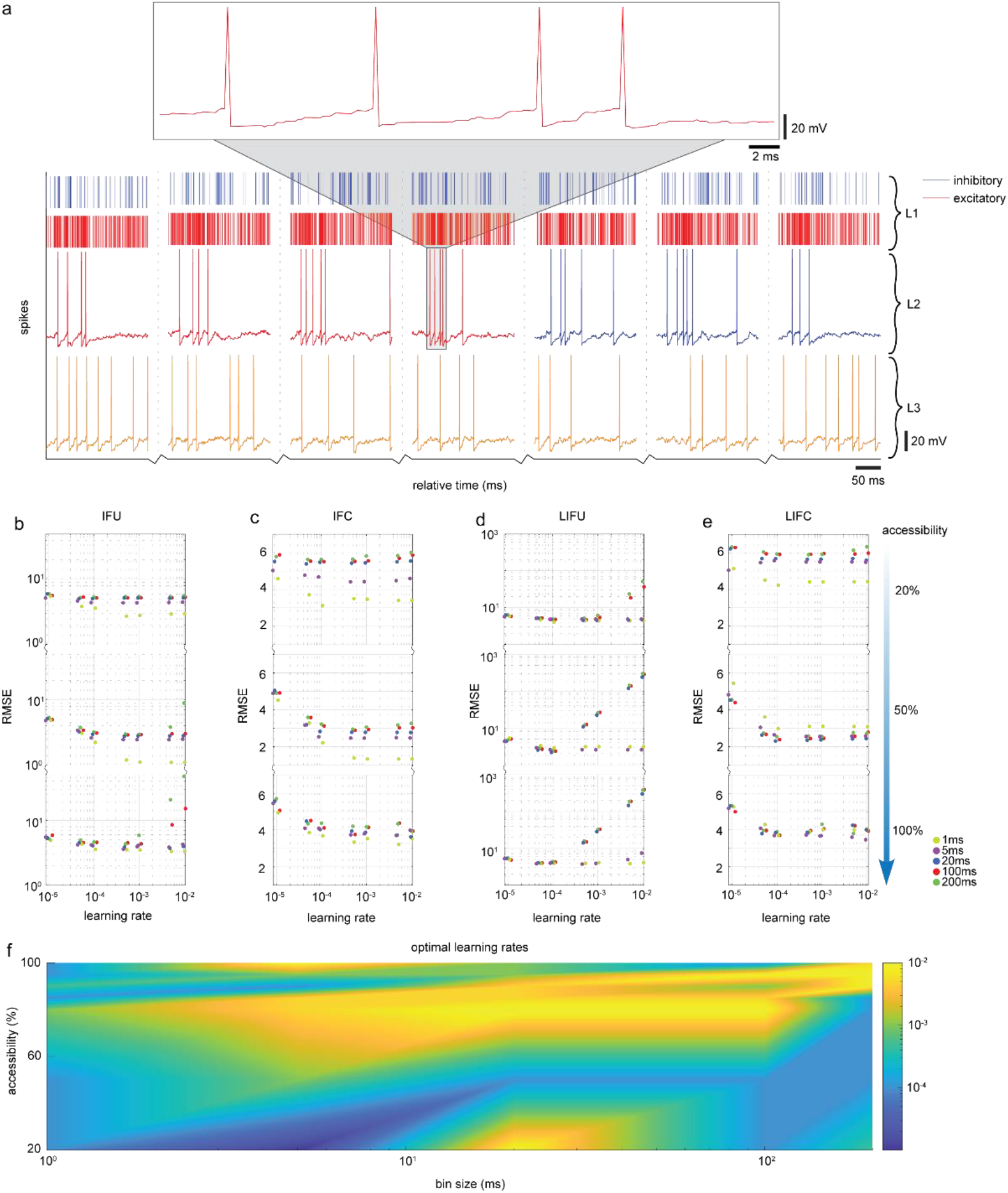
Reconstructing connectivity from firing rates of bursting network subpopulations. **(a)** Demonstration of layer 2 (L2) bursting behavior resulting from superimposed layer 1 (L1) spike trains and corresponding layer 3 (L3) voltage trace. L2 neurons shown have strong excitatory and inhibitory connections to L3 neuron (excitatory - left; inhibitory - right). **(b-e)** RMSE results for IFU, IFC, LIFU and LIFC under varying L2 accessibility conditions. **(f)** A map is provided for determining optimal learning rates for given accessibility and bin size under LIFC conditions.

## 4. DISCUSSION

This study delineates a recipe for inferring neural connectivity based on supervised learning of temporally blurry synaptic data. By adapting the perceptron model to low temporal resolution scenarios, we demonstrate optimized reconstruction of feedforward presynaptic connections based on correlation between performance, learning rate, temporal resolution of the input data, and the underlying presynaptic firing rates. Furthermore, we show that properly inferred connections can be applied to predict network activity at the single-spike level with relatively high precision, with implications for improving reverse engineering of neuronal network activity using constrained recording technologies.

Our protocol prescribes optimal learning rates to decode connectivity of networks receiving low sampling rate signals with no assumptions made to spiking patterns within a given time bin. We test this generalized method on networks displaying physiological spike bursting patterns and find optimized learning rates for reconstructing presynaptic weights. Future adaptations can be tailored to train over such specialized spiking patterns (42), including spikes phase locked to sensory stimulus (43), network oscillations, and log-normal spiking behavior generally seen in the brain (4). By redistributing single spikes within a given temporal bin during learning iterations, our perceptron variant can optimize performance preferentially to neural activity of a specific persuasion and can potentially achieve more accurate connectivity maps and single-spike predictions with biological data as input. Assumptions on spiking behavior can be integrated with approaches shown to successfully reconstruct undersampled networks with only sparse datasets observable from subpopulations of the network (34), and can in turn allow for processing of larger networks and datasets commonly reconstructed using generalized linear models that overcome computationally exhaustive data processing (36,44). Other relevant postprocessing tools used to deconvolve calcium readouts, EEG signals and similar temporally limited readouts into time-resolved action potentials (17,18,34,45) can also improve the ability to decipher connectivity by serving as precursors for upgrading the input data prior to learning of weights. Additionally, single-neuron calcium spikes and similar biophysical events have specialized nonlinear features that are leveraged by deconvolution methods to decode underlying voltage spikes by supervised learning of large amounts of data (18,19). Future variations of the method presented here will mimic specific signal properties relevant to biophysical processes and train over less naïve presynaptic input and signal shapes, as well as use leaky integrate and fire neurons more suitable for recreating tonic firing and bursting network activity (46,47).

Comprehensive reconstruction of mammalian cortical and subcortical microcircuitry uses ultrastructural, morphological, functional and computational analyses to reveal diverse connectivity that can reenact brain states in silico (48). Here, we presented a proof-of-concept study for naive, linear feedforward networks with uniform distribution of presynaptic connectivity. Processing of diverse biological datasets to explain realistic networks will rely on applying the modified perceptron to recurrent neuronal circuits with interconnected hubs and heterogeneous weight distributions comprising both unidirectional and reciprocal connections (48–50). Furthermore, recordings occurring at time scales involving both short-term and long-term plastic processes related to learning and memory will require dynamic assignment of weights. In order to account for these and other nonlinear functional and topological network traits, dynamically responsive approaches where learning rates are not held constant during iterative training are expected to allow for improved performance (51). Our training sets correspond to neurobiological context, with millisecond timescales, all-or-nothing membrane potential fluctuations frequencies, and typical synaptic strength, but can be used more broadly for reverse engineering functional biological networks limited by temporally constrained readouts.

## Acknowledgments

This research was funded by NIH grant K01EB027184 and DP2NS122605 to AH. ADV performed the research. ADV and AH wrote the manuscript. XR and JE participated in coding and data analyses. We thank Dr. Etay Hay for supplying reference code and useful comments on the manuscript.

